# Detecting sources of immune activation and viral rebound in HIV infection

**DOI:** 10.1101/2022.06.08.495415

**Authors:** Stephen W. Wietgrefe, Lijie Duan, Jodi Anderson, Guillermo Marqués, Mark Sanders, Nathan W. Cummins, Andrew D. Badley, Curtis Dobrowolski, Jonathan Karn, Amélie Pagliuzza, Nicolas Chomont, Gérémy Sannier, Mathieu Dubé, Daniel E. Kaufmann, Paul Zuck, Guoxin Wu, Bonnie J Howell, Cavan Reilly, Alon Herschhorn, Timothy W. Schacker, Ashley T. Haase

## Abstract

Antiretroviral therapy (ART) generally suppresses HIV replication to undetectable levels in peripheral blood, but immune activation associated with increased morbidity and mortality is sustained during ART, and infection rebounds when treatment is interrupted. To identify drivers of immune activation and potential sources of viral rebound, we modified RNAscope in situ hybridization to visualize HIV-virus producing cells as a standard to compare the following assays of potential sources of immune activation and virus rebound following treatment interruption: 1) EDITS (envelope detection by induced transcription-based sequencing) assay; 2) HIV-Flow; and 3) Flow-FISH assays that can scan tissues and cell suspensions to detect rare cells expressing env mRNA, gag mRNA/Gag protein and p24 respectively; and 4) an ultrasensitive immunoassay that detects p24 in cell/tissue lysates at subfemtomolar levels. We show that the sensitivity of these assays is sufficient to detect a rare HIV-producing/env mRNA+/p24+ cell in a million uninfected cells. These high-throughput technologies thus provide contemporary tools to detect and characterize rare cells producing virus and viral antigens as potential sources of immune activation and viral rebound.

**Importance:** Anti-retroviral therapy (ART) has greatly improved the quality and length of life for people living with HIV, but immune activation does not normalize during ART, and persistent immune activation has been linked to increased morbidity and mortality. We report a comparison of assays of two potential sources of immune activation during ART: rare cells producing HIV virus or the virus’ major viral protein, p24, benchmarked on a cell model of active and latent infections and a method to visualize HIV-producing cells. We show that assays of HIV Envelope mRNA (EDITS assay) and gag mRNA and p24 (Flow-FISH, HIV-Flow and ultrasensitive p24 immunoassay) detect HIV-producing cells and p24 at sensitivities of one infected cell in a million uninfected cells, thus providing validated tools to explore sources of immune activation during ART in the lymphoid and other tissue reservoirs.

## Introduction

ART has greatly improved the quality and length of life for people living with HIV (PLWH), but treatment must be continued indefinitely, because infection quickly rebounds from persistent viral reservoirs if treatment is interrupted [1,2]. In addition to the challenges that recrudescent infection poses for curing HIV infection, immune activation (IA) decreases but does not normalize during ART, and persistent IA has been linked to increased morbidity and early death from inflammatory mediated conditions [3–10]. Thus, it will be important to identify the sources of residual IA to devise mitigating strategies to improve long-term health outcomes for treatment of people PLWH.

Here we report a comparison of assays of cells that harbor potentially replication competent proviruses or defective proviruses that could nonetheless generate viral antigens as potential drivers of IA: 1) EDITS [11] assay to detect cells expressing env RNA; 2) HIV FISH/Flow [12] to detect cells expressing gag RNA and p24; 3) HIV-Flow [13] to detect cells expressingp24; 4) p24 ultrasensitive immunoassay [14] to detect p24 at subfemtomolar levels. Because env+ and p24+ cells and p24 antigen during ART are expected to be rare and in small quantity, we further asked whether the limits of detection (LOD) for the EDITS, HIV-Flow, HIV-FISH/Flow, and ultrasensitive p24 high throughput assays would respectively be able to detect rare env RNA+, gag mRNA/p24+ or p24+ cells in a background of a million or more uninfected cells.

We developed a method to visualize HIV-producing ACH-2 cells to serve as a standard to evaluate these high throughput assays for two reasons. First, as a latently infected cell line in which virus production can be reactivated by treatment with phorbol esters or TNF-α, ACH-2 cells as a surrogate for reactivated latently infected cells [15]. Second, we knew from previous ISH studies [16] that reactivation of ACH-2 cells results in levels of HIV RNAs, Gag and Env protein that should be detectable in the EDITS and p24 assays and thus serve as standards to evaluate the sensitivity of these approaches to detect rare cells with HIV RNAs and antigen.

## Results

### RNAscope ISH to detect HIV-producing cells

The method we originally developed to detect SIV-producing cells [17] amplified the signal from SIV RNA in virions by tyramide signal amplification to deposit sufficient - product under diffusion limiting conditions to reveal visible virions at the resolution of visible or fluorescent light microscopy. To reveal HIV-producing cells, we modified a contemporary RNAscope ISH protocol [18] to visualize not only intracellular HIV RNA but also HIV RNA in virions associated with the infected cells.

The substitution of the ELF 97 alkaline phosphatase substrate in RNAscope ISH deposited sufficient ELF-97 product around HIV RNA in virions so that they were clearly visible in ACH-2 cells (Fig 1). Prior to induction, about 5 percent of ACH-2 cells undergoing spontaneous reactivation score as HIV-producing cells. Virus-producing cells were not detected in the remaining 95 percent prior to induction, in CEM or Jurkat HIV-negative cells, or in induced ACH-2 cells with plus sense probes (not shown). Following induction, 100 percent of the cells were visibly producing HIV virions. In counts of 78 cells where the z-series 3D images included all or nearly all the cells, the average virion count per induced ACH-2 cell was 558, SD, 66. The average size of virions rendered visible by the deposition of the ELF-97 substrate was ~ 260 nm, consistent with the diffraction limit for ELF 97 emission at 530 nm. Thus, the in situ single cell assay detects HIV-producing ACH-2 cells at the resolution of immunofluorescence microscopy, and documents virus production in all induced cells, thus providing enabling technology to visualize HIV producing cells as a standard for subsequent assay comparisons.

**Fig 1.**
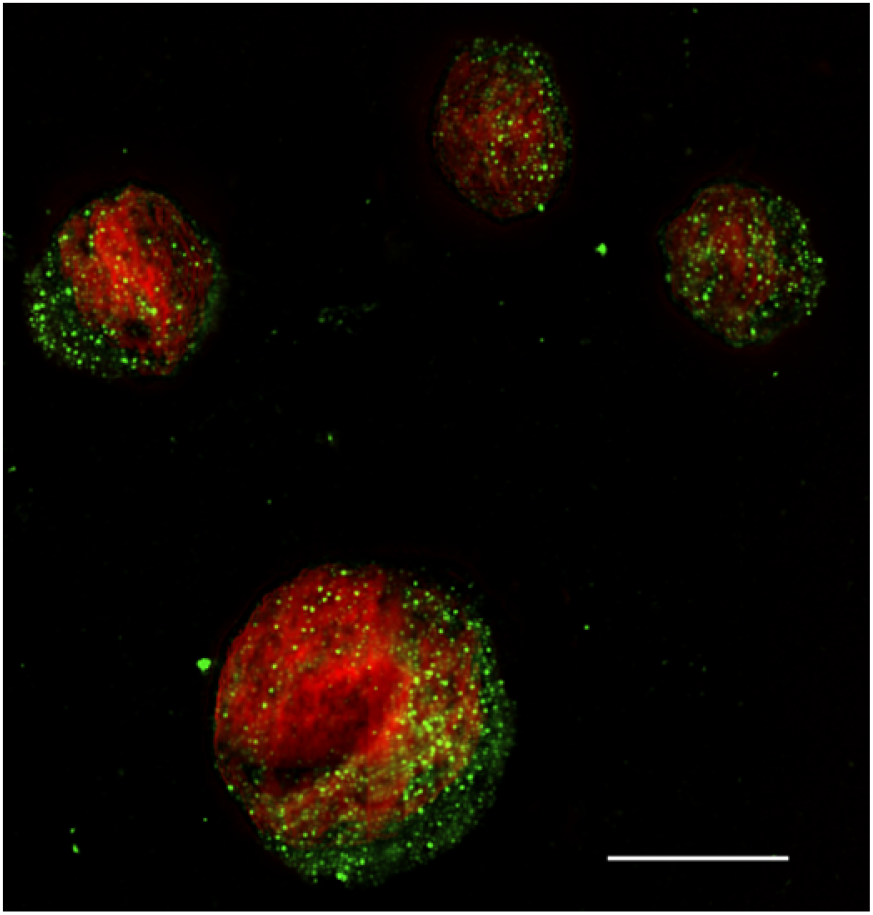
RNAscope ISH/ELF 97 detection of HIV producing cells in 5-μm sections of fixed and embedded induced ACH-2 cells. HIV virions appear green in the image. Nuclei counter-stained red with TOTO 3. In this 2D image, the largest number of virions are associated with a whole cell in the focal plane of the z-series. Scale bar= 10 μm.

### Relationship between HIV-producing cells and production of infectious virus

Before evaluating the sensitivity of the high throughput assays with HIV producing ACH-2 cells as the standard for comparison, we also investigated the relationship between visible production of virions and production of infectious virus. For this purpose, we used a previously described modified viral outgrowth assay (QVOA) [19,20] to document production of infectious HIV by induced ACH-2 cells and determine how accurately the QVOA would estimate the frequency of cells with inducible replication competent proviruses. Serial dilutions of mixed ACH-2 cells and Jurkat T cell samples were induced by co-culture with irradiated, CD8-depleted allogeneic PBMC feeder cells plus anti-CD3 and IL-2. The induced cells were co-cultured with anti-CD3 stimulated, CD8-depleted PBMC target cells for 14 days at which time infectious units per million cells (IUPM) were estimated by limiting dilution statistics on p24+ cultures.

The QVOA documented production of infectious virus by induced ACH-2 cells, albeit at 28 to 81 percent of the expected number of virus-producing cells in samples with 1, 10 and 100 infectious units per million (IUPM) cells (Table 1). We attribute the lower estimates to the well-documented underestimates of the frequency of latently infected cells harboring replication competent intact proviruses [21,22] because of the inability at least in part to amplify infection from small numbers of cells to detect all the cells with a replication competent provirus in QVOA assays.

**Table 1.**
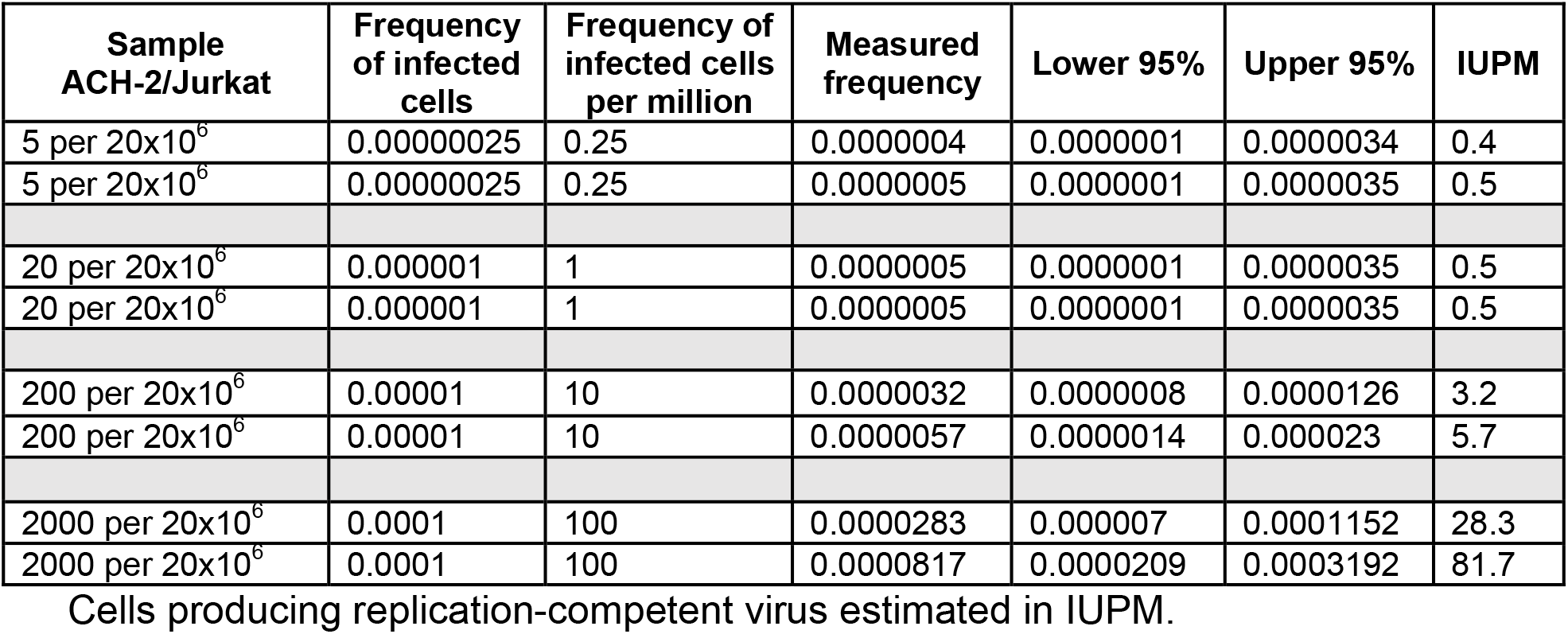
QVOA assay of ACH-2 cells.

### EDITS assay

The EDITS assay measures spliced env mRNA by next generation sequencing of the major HIV-1 env RNA splice junction to detect the late spliced viral transcript as a measure that like the intact proviral DNA assay [23], can distinguish between replication competent and defective proviruses. We evaluated concordance between the EDITS assay and virus producing cells in induced ACH-2 cells diluted with HIV-negative CEM cells to cover a range of 2.5 to 2500 ACH-2 cell equivalents in 1.25 × 10^6^ cells in the typical format for this assay. Total RNA isolated from un-stimulated or ACH-2 cells induced with PMA was converted to cDNA for PCR amplification of a PCR product with the major env splice junction for ion torrent sequencing. Reads mapping to a synthetically spliced HXB2 sequence were scored and expressed as an equivalent number of cells harboring HIV-1 per 10^6^ cells, using a standard curve determined for activated primary memory CD4 T cells infected with replication-competent HIV-1 carrying a GFP reporter. In the linear range of the assay [11] in samples with 2.5 to 160 induced ACH-2 cells, the EDITS assay results were highly correlated but consistently slightly higher compared to the number of virus-producing ACH-2 cells expected after stimulation (Table 2 and Fig 2), an overestimate consistent with induction of env mRNA without virion production in ~ 15 to 30% of the cells. In the samples with 320 and 2500 ACH-2 cells outside the linear range, the EDITS assay underestimated the number of virus-producing cells respectively by about 15 and 80 percent.

**Table 2.** EDITS assay in triplicate of ACH-2 cells induced with CD3/CD28 T cell activator in the numbers shown.

| Induced ACH-2 per<br>1.25 x10<sup>6</sup> CEM Cells | EDITS Estimated<br>Stimulated Cell Equivalents |  |  |
| --- | --- | --- | --- |
| 2.5 | 2.54 | 3.35 | 2.95 |
| 5 | 5.64 | 6.45 | 6.04 |
| 10 | 11.5 | 10.6 | 11.1 |
| 20 | 24.9 | 28.5 | 26.7 |
| 40 | 53.0 | 53.6 | 53.3 |
| 80 | 83.4 | 91.4 | 87.4 |
| 160 | 183 | 189 | 186 |
| 320 | 273 | 278 | 276 |
| 2500 | 456 | 446 | 451 |

**Fig 2.**
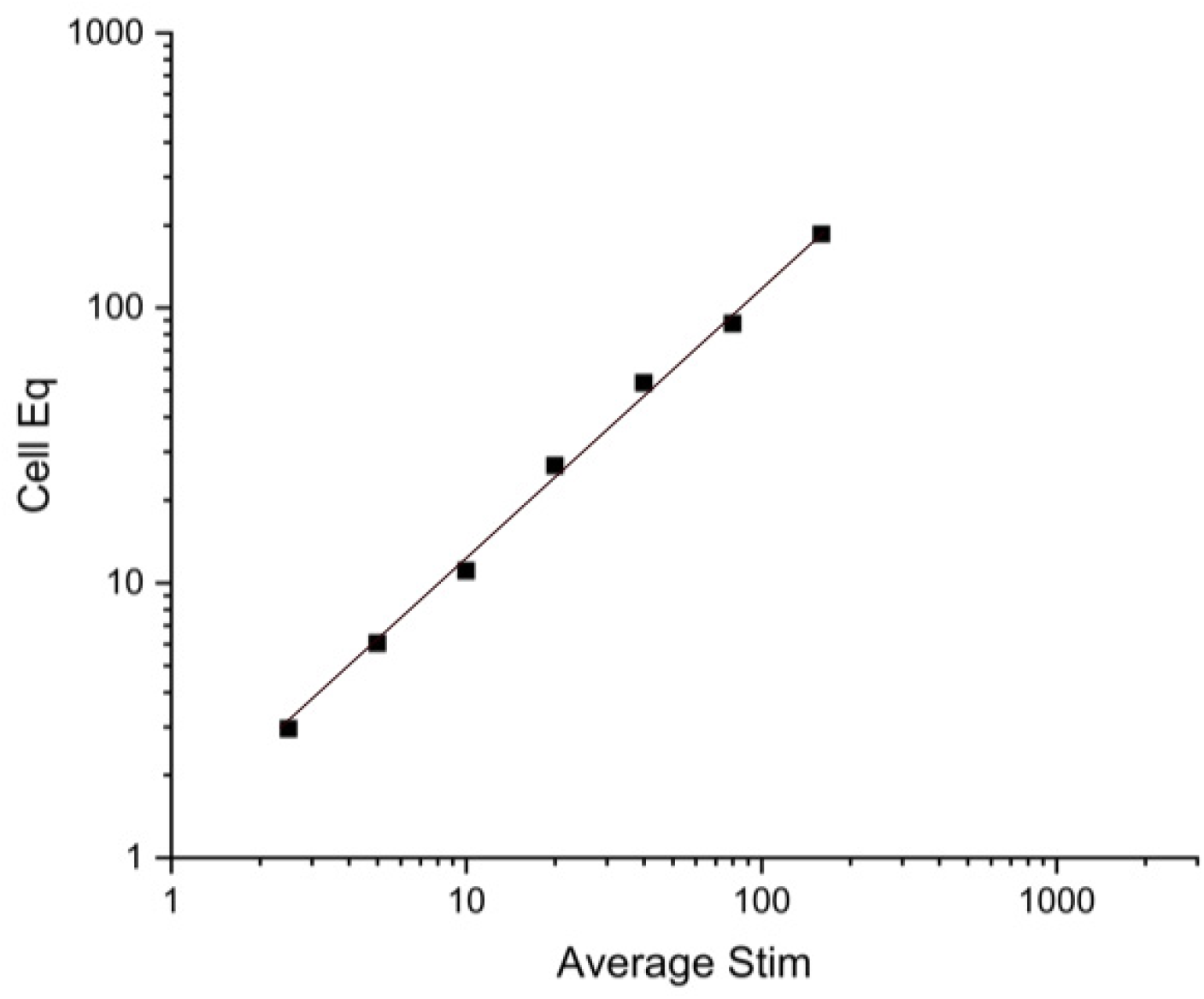
Logarithmic plot of average of replicates of induced ACH-2 cell equivalents (Eq) and EDITS assay results in Table 2. Linear relationship to 160 cells. Pearson’s r=0.99734.

### HIV-Flow Assay

The HIV-Flow Assay measures the frequency of cells with translation-competent HIV proviruses that produce detectable levels of p24 [34]. In this flow cytometry-based assay, p24+ cells are identified by combining two monoclonal antibodies to p24 (KC57 and 28B7) coupled to two different fluorochromes, thereby improving the specificity of the measurement. We determined the relative efficiency of the HIV-Flow assay for HIV-producing/p24+ cells in a dilution series of 1, 10, 100 and 1000 induced ACH-2 cells in a million CEM. As shown in Table 3 and Fig 3, HIV-Flow gave frequencies highly correlated with but consistently slightly higher than the expected values, possibly reflecting detection of uninfected cells to which released virions had attached.

**Table 3.** HIV-flow assay dilution series.

| Expected Frequency<br>HIV positive cells/$10^6$ cells | Measured Frequency<br>p24+ cells/$10^6$ cells |
| --- | --- |
| 1000 | 1068 |
| 128 | 152 |
| 64 | 83 |
| 32 | 47 |
| 16 | 18 |
| 8 | 9 |
| 4 | 6 |
| 2 | 5 |
| 1 | 2 |

Expected versus determined frequency of induced ACH-2 cells per $10^6$ cells.

**Fig 3.**
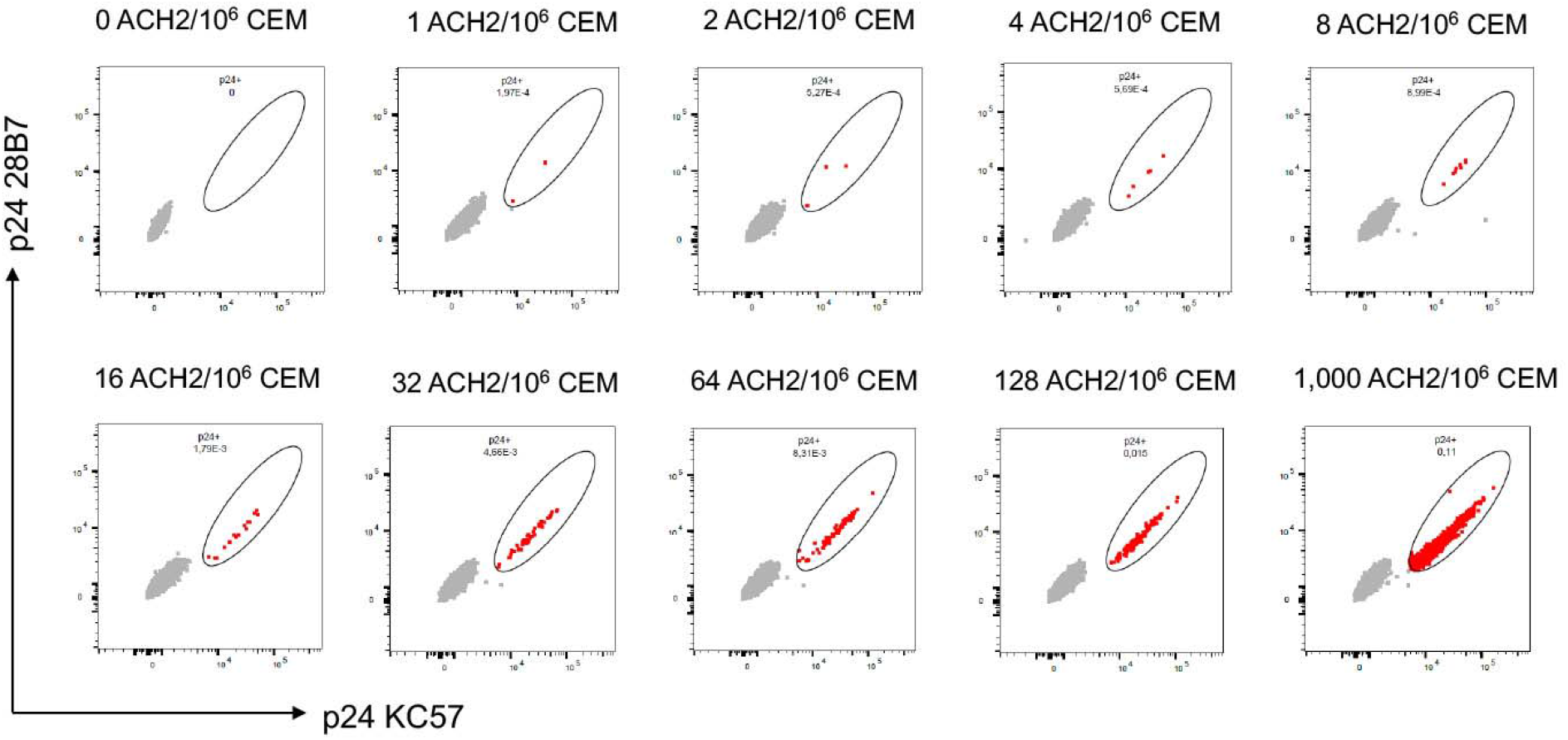
(A) Flow cytometry dot plots showing the frequency of induced ACH2 cells serially diluted in CEM cells measured by HIV-Flow. The expected frequencies are indicated at the top of each dot plot. p24+ cells are depicted in red, p24-cells in grey.

**Fig 3.**
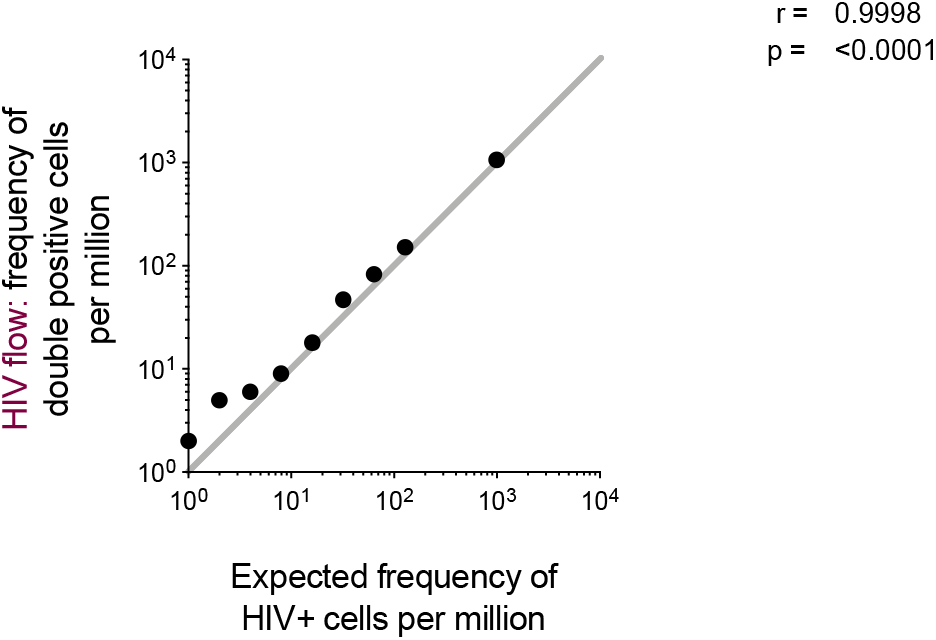
(B) Correlation between HIV-flow estimates of p24+ cells per million cells and the expected frequency in the serial dilution series.

### HIV RNA Flow-FISH assay

The HIV^RNA/Gag^ assay is a RNA-flow cytometric fluorescent *in situ* hybridization (RNA Flow-FISH) technique that combines detection of HIV mRNAs and Gag protein (p24) staining [35] with a sensitivity of detection of 0.5-1 double positive CD4+ T cells per million uninfected CD4+ T cells. Concurrent detection of HIV transcription and translation products enables distinction between translation-competent cells and translation-incompetent cells. To assess the specificity and linearity of the HIV^RNA/Gag^ assay, we spiked reactivated latently infected ACH2 cells into uninfected CEMx174 cells. In the absence of reactivation, HIV^+^ events, defined as cells co-expressing *gag* mRNA and Gag protein (HIV^RNA+/Gag+^) were detected in 5.9% of ACH2 cells (Fig 4A), in good agreement with the RNAscope ISH analysis cited above for un-induced ACH-2 cells. Time-course experiments showed that a 24-hour stimulation with PMA/Ionomycin induced high expression of Gag products in ACH-2 cells (95.2% HIV^RNA+/Gag+^ cells) (Fig 4A), and this condition was therefore selected for the spiking experiments. The input corresponding to the highest frequency of reactivated ACH2 into the CEMx174 line was measured by flow cytometry at 1500 HIV+ events per million cells (Fig 4B), consistent with the initial input planned (see Methods), while the false positive event rate in pure CEMx174 cells was low (1 HIV^RNA+/Gag+^ cell detected in 700,000 cells) (Fig 4C). Spiking dilutions ranged from a theoretical expected frequency of 1.5 to 1500 events per million cells (Fig 4D). The experimental frequencies of HIV+ events experimentally identified by HIV mRNA/protein co-staining showed excellent linearity and consistency down to the lowest dilution tested, except at the two lowest spiking dilutions (<6 expected events/million), for which the variability noted between the observed and expected numbers of HIV+ events can be reduced by analyzing larger numbers of cells [24].

**Fig 4.**
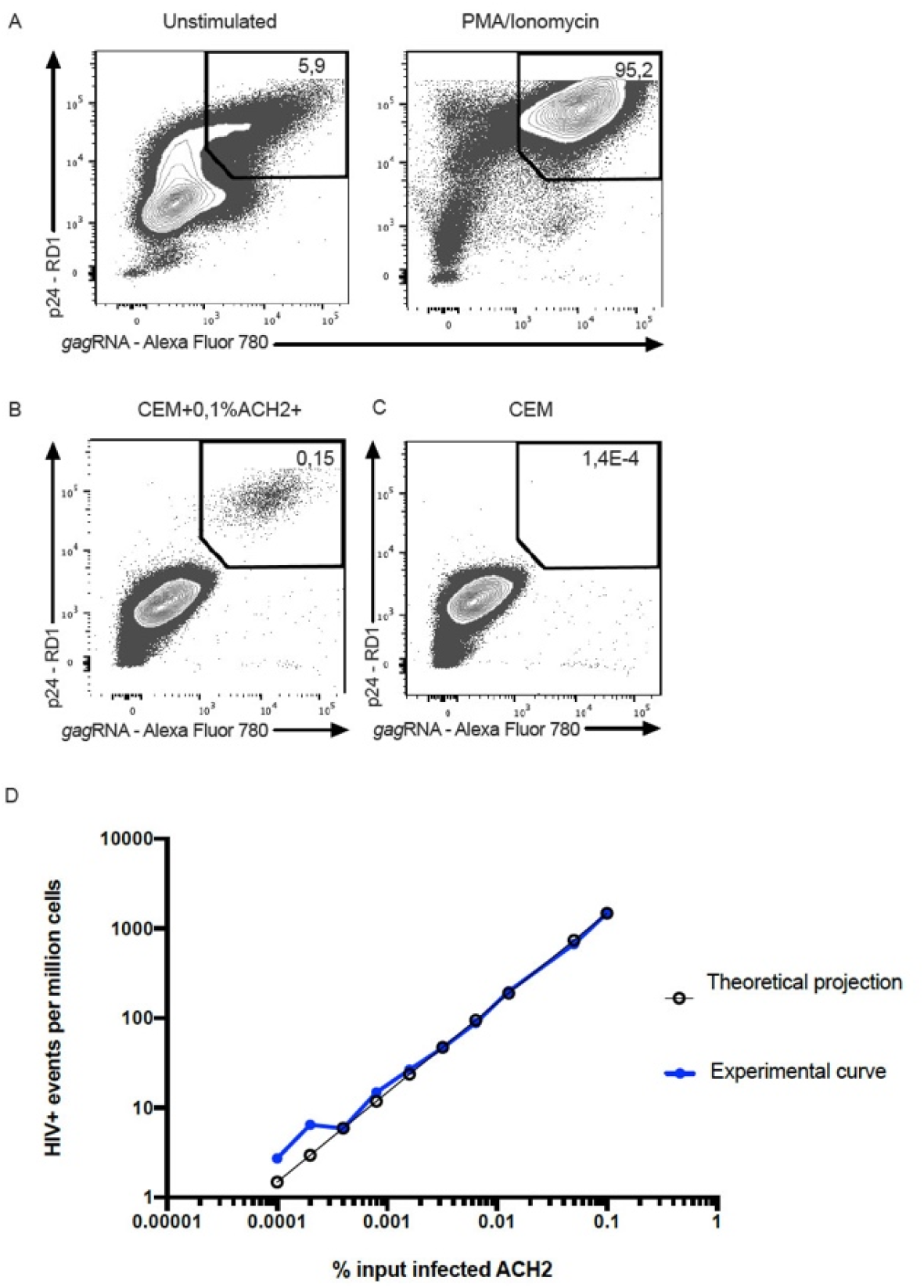
Flow cytometry example plots of the HIV RNA Flow-FISH assay and correlation between measured and expected frequencies of infected cells in spiking experiments. (A) p24+ gag mRNA+ ACH-2 cells before and after induction with PMA/Ionomycin. (B) Highest frequency of reactivated ACH2 spiked into the CEMx174 line. (C) False positive event rate in pure CEMx174 cells. (D) Theoretical expected frequency of 1,5 to 1,500 events per million cells and experimental frequencies of HIV+ events experimentally identified by HIV mRNA/protein co-staining.

### HIV ultrasensitive p24 immunoassay

Digitized immunoassays using single-molecule array (SIMOA®) technology have extended the sensitivity of detection of HIV p24 into the femtomolar range and we and others have reported further enhancement of the technology to achieve sensitivity limits of detecting sub-femtomolar levels of HIV p24+ protein [14,25–28]. Here we investigated the sensitivity of detecting p24+ induced ACH-2 cells in a matrix of a million uninfected PBMCs with our ultrasensitive assay, and show that the ultrasensitive p24 assay detects a single p24+ cell in a million PBMCs (Table 4). Because of the variability between triplicate samples that we attribute to clumping of the induced ACH-2 cells, we tested lysates of induced ACH-2 cells diluted into PBMCs corresponding to samples containing 1 or 10 induced ACH-2 cells, and again detect a single positive cell per million. There was good agreement between samples with a CV <10%, and the expected 10-fold higher levels of p24 in the samples with 10 induced ACH-2 cells per million PBMCs.

**Table 4.** Detection of p24 in triplicate samples of induced ACH-2 cells diluted into 10^6^ uninfected PBMCs.

| Induced ACH-2 Cells per Million PBMCs | p24 #1 | p24 #2 | p24 #3 | Average p24 Concentration pg/mL | Std. | CV (%) |
| --- | --- | --- | --- | --- | --- | --- |
| 0 | 0.031 | 0.012 | 0.027 | 0.023 | 0.01 |  |
| 1 | 2.928 | 0.222 | 0.291 | 1.147 | 1.54 |  |
| 10 | 24.984 | 12.166 | 13.585 | 16.192 | 7.03 |  |
| 100 | 106.574 | 121.748 |  | 114.161 | 10.73 |  |
| 1 | 1.49 | 1.36 | 1.59 | 1.478 | 0.12 | 7.8 |
| 10 | 14.45 | 14.74 | 15.36 | 14.853 | 0.46 | 3.1 |

Because of the variability between triplicate samples above the heavier line, attributed to clumping of cells, the samples below the line are direct lysates containing 1 or 10 induced ACH-2 cells diluted in a million PBMCs.

## Discussion

We describe and compare here a suite of tools to detect env and gag mRNA or p24-expressing cells in high throughput scans of rare cells in a background of a million or more uninfected cells. These technologies provide validated standards and approaches to quantifying HIV reservoirs that are highly relevant to identifying sources of virus before, during and off ART as well as drivers of IA that can guide the design of strategies to potentially prolong remissions off ART, and mitigate IA for longer healthier lifespans for PLWH.

The EDITS, HIV-Flow, HIV-FISH-Flow and ultrasensitive p24 assays proved to be facile, rapid, sensitive, and accurate in detecting respectively single cells with env or gag mRNA and p24 in a million uninfected cells. These high throughput approaches thus can provide a quick read on the likelihood that peripheral blood or tissue samples harbor cells with respectively sufficiently intact proviruses that can be reactivated to produce env mRNA or translationally competent proviruses that generate p24.

The EDITS assay primers and amplicons to detect late spliced env mRNA overlap the packaging site amplicon in the IPDA assay [23], and both assays generate sequences from the 3’ end of proviral genomes (Fig 5). Moreover, generating env mRNA requires functional Tat and Rev, consistent with the possibility that the EDITS assay env transcript might score for largely intact functional genomes as in the IPDA assay. We show six examples of deleted proviruses (Fig5B) to demonstrate how the EDITS and IPDA assays might score each virus and found that both assays correctly score proviruses a, c-e as proviruses with deletions. The EDITS assay misses the small deletion in gag-pol in provirus b detected in the IPDA assay, and both assays incorrectly score provirus f as intact, based solely on the sequences in the amplicons. However, provirus f would likely lack Tat and Rev, and would therefore be correctly scored as impaired in the EDITS assay. We think this comparison suggests that the two assays should generally be in good agreement, and this expectation has been satisfied experimentally in data on the impact of the IL-15 superagonist N-803 on the frequency of inducible HIV provirus in peripheral blood mononuclear cells [29].

**Fig. 5.**
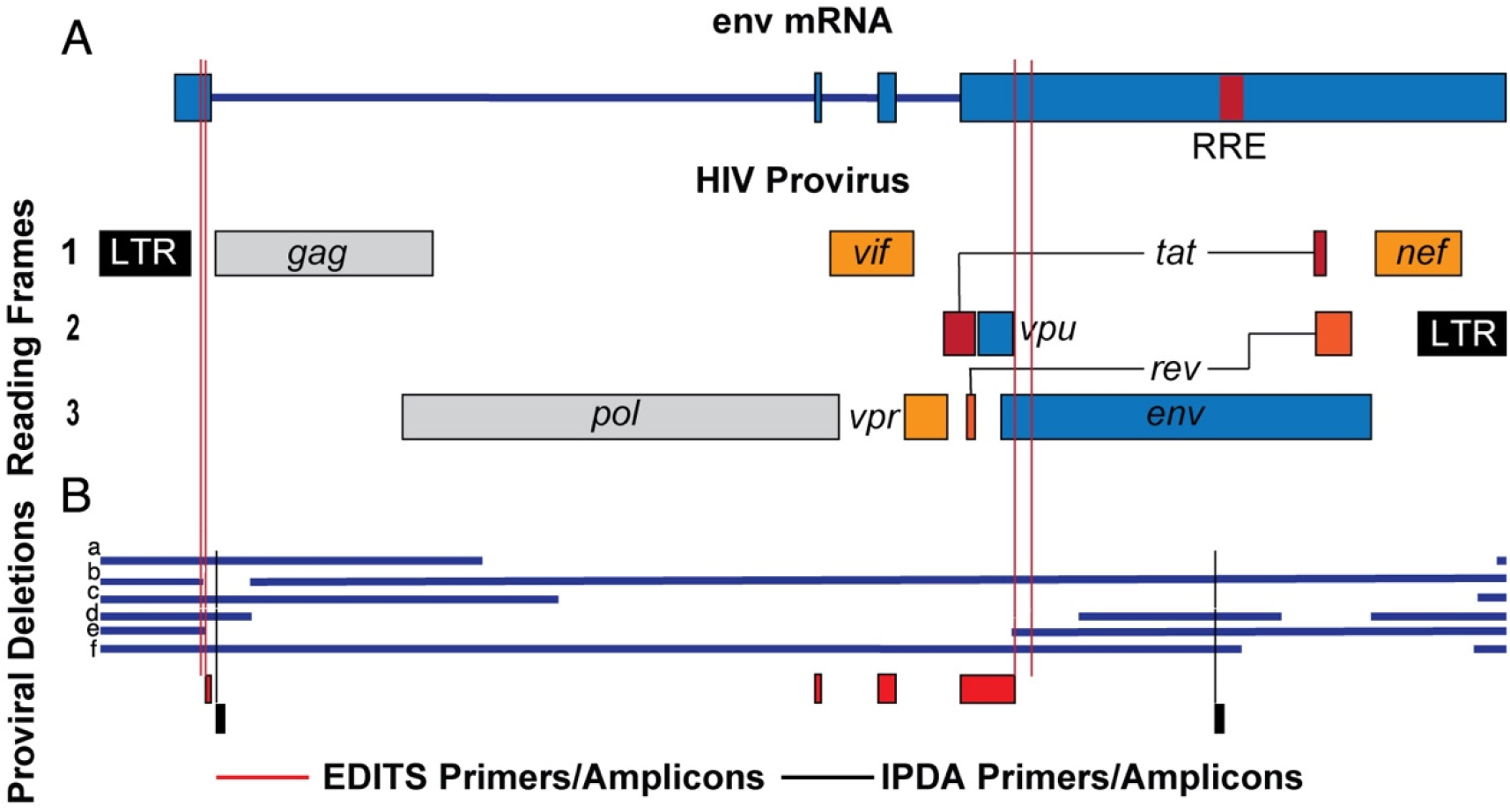
Comparison of EDITS and IPDA assays for identifying HIV proviruses with deletions. (A) Schematic of HIV provirus and env mRNA. Red lines indicate positions of the EDITS assay primers. (B). Comparison of the assays to detect deletions in proviruses a-f. EDITS primers and amplicons, red; IPDA primers and amplicons, black.

The relationship of HIV-p24+ cells detected in the p24 assays to HIV virus producing cells is indeterminate. However, we can conclude that detection of p24+ cells at the least is potentially relevant to sustaining IA and identifying cells that could be targeted by HIV-specific CD8+ T cells. Collectively, the high throughput approaches described here provide well-characterized tools to explore mechanisms of HIV persistence and IA during ART and to devise rational strategies to sustain remissions off ART and mitigate IA.

## Materials and Methods

### ACH-2 cells

ACH-2 cells, and uninfected CEM or Jurkat cells were grown in RPMI with 10% heat inactivated FCS, glutamine, and 1X antibiotic-antimycotic (FISHER). ACH-2 cells were activated in some experiments by growth with 50 ng/ml PMA for 24 hrs. Cells were collected by centrifugation at 450g x 7min. and at least two washes in PBS, 0.5% BSA, 2mM EDTA. Cell mixtures were created by suspending uninfected cell lines in 1/10 culture volume of buffer, ACH-2 cells in their culture volume, and counting cells with trypan blue staining in a Countess II automated cell counter in duplicates. CEM or Jurkat cells were then adjusted to 10^7^ cells/ml with buffer, and ACH2 cells (induced or uninduced) to 10^5^ or 10^6^ cells/ml. An initial cell mixture of 10^3^ ACH-2 cells in 10^7^ uninfected cells/ml was created and diluted serially 2X or 10X with 10^7^ uninfected cells/ml. Cell mixtures were frozen, extracted or fixed as required by the specific assay, per published assay protocols.

### Detection and characterization of HIV producing cells by RNAscope ISH combined with IFA

We varied conditions in our previously published RNAscope ISH method [18] to be able to not only detect HIV intracellular RNA but also clearly reveal virions associated with the cells. The critical changes in the published protocol were: 1) reduced time sections were boiled in RNAscope Pretreat citrate buffer to 10 minutes; 2) sections incubated for 20 minutes at 40°C in a more dilute (1:10) Pretreat 3 reagent 3 (protease digestion solution, 2.5 μg/ml); and 3) following AMPs 1-6 applied from the RNAscope 2.5 HD Detection Reagent Red kit (Advanced Cell Diagnostics), ten to thirty minute room temperature incubation with ELF 97 Phosphatase substrate. For the images of HIV-1 production by induced ACH-2 cells shown in Fig 1, nuclei were stained with To-Pro-3 (Thermo Fisher Scientific).

### Confocal plus deconvolution imaging

Fluorescent confocal images of ELF 97 and To-Pro 3 labeled cells were acquired in a Nikon Ti-E inverted microscope equipped with a CFI SR Apo 100X TIRF Oil Immersion Objective Lens NA 1.49 and a Nikon A1R confocal scan head controlled by Nikon Elements software (5.11). ELF97 excitation was provided by a 403nm laser and emission collected between 500-550 nm. To-Pro 3 excitation was provided by a 640nm laser and emission collected between 663-738 nm. The confocal aperture was set to the minimum value of 14 μm (0.2AU). The images were oversampled in X, Y (41 nm pixel). A 200 nm Z step was used in three-dimensional acquisitions and images were subjected to automatic iterative deconvolution with Nikon Elements 5.20. Analysis was done in Elements ver. 5.20 using the GA3 pipeline including 3D object measurement and FIJI software using find maxima after making maximum intensity projections of rolling ball background corrected imaging for particle counting. Channels were pseudo-colored and merged. EDF projections of the z-stacks are presented. University of Minnesota-University Imaging Centers, http://uic.umn.edu/.

### QVOA

Quantitative viral outgrowth assays were performed as previously described [20]. Briefly, six 5-fold serial dilutions of mixed ACH-2 cells and Jurkat T cell samples were plated in duplicate starting at 1×10^6^ cells per well. These samples were co-cultured with irradiated, CD8-depleted PBMC feeder cells, and stimulated with anti-CD3 (OKT3, 1 μg/mL) and IL-2 (50 IU/mL) overnight. On day 1 and day 7 post stimulation, 1×10^6^ anti-CD3 stimulated, CD8-depleted PBMC target cells were added to each well. Supernatant p24 was measured on day 14 by ELISA (Zeptometrix) per manufacturer’s protocol. Individual wells were determined to be positive or negative for p24 based on manufacturer determined cut-off values. Infectious units per million were estimated by limiting dilution statistics using L-Calc™ Limiting Dilution Software (STEMCELL Technologies).

### EDITS assay

The frequency of env mRNA+ induced ACH-2 cells was determined by the EDITS assay, as described by Das et al [11]. In brief, ACH-2 cells were induced with 50 ng/ml PMA for 24 hrs. Total RNA was isolated using the Qiagen RNeasy purification system (Qiagen, 74134) following the manufacturer’s protocol. The entire sample was used as template in a one-step RT-PCR reaction (Thermo Scientific, AB-4104A). After cDNA synthesis and PCR, 2 μl of the reaction was used as template for a subsequent round of nested PCR using a high fidelity Phusion Flash polymerase (Thermo Scientific, F548). Primers were designed to bind to either side of the HIV Env RNA splice junction using highly conserved regions of HIV. For the first-round PCR, the forward primer was located at position 570-591 and the reverse primer was at positions c6442 and c6426. In addition to the priming sequence, the reverse primer has a synthetic GEX R-AATGATACGGCGACCACC sequence placed directly after the priming region to allow for further amplification using nested PCR. The RT-PCR product was further amplified by nested PCR with a nested forward primer at position 610-631 and a nested reverse primer at c6324-6345. To allow for NGS sequencing, Ion torrent A forward and Trp reverse adapters were added to the nested primer sets, as well as a unique barcode in the forward primer, to allow for multiplexing of samples. Samples were then pooled and primers were removed using GeneJET NGS cleanup kit (Fisher Scientific, FERK0852). DNA concentrations were measured by a Qubit fluorescent reader and 300 pg of the pooled sample was then sequenced using an Ion Torrent Sequencing system following the manufacture’s protocol. Barcodes were separated by sample using the Ion Torrent Browser and all reads were filtered to remove short products (under 80 bp) and only reads that contained the GEX reverse sequence were retained. The filtered reads were then mapped to a synthetically spliced HXB2 sequence, and total mapped reads were scored. The number of mapped reads was then converted into the equivalent number of cells harboring HIV-1 per 10^6^ cells using a standard curve generated from activated memory cells infected with replication-competent HIV-1–GFP virus, and sorted by flow cytometry into single wells of a 96-well plate. Samples for the standard curve contained between 1 and 300 infected cells per well and 1.25 × 10^6^ uninfected cells.

### HIV-Flow assay

The frequency of p24+ cells was determined as previously described [12]. Briefly, nine serial dilutions of ACH2 cells in CEM cells were stimulated with 162nM PMA (Sigma, P8139) and with 1 μg/mL ionomycin (Sigma, I9657). After 24h of stimulation, cells were collected, resuspended in PBS and stained with the Aqua Live/Dead staining kit for 30min at 4°C. Cells were stained in PBS +4% human serum for 30min at 4°C (CD3 A700, CD4 BUV496, CD8 BUV395). The fixation/permeabilization step was performed with the FoxP3 Transcription Factor Staining Buffer Set (eBioscience, 00-5523-00) following the manufacturer’s instructions. Cells were then stained with anti-p24 KC57 (R&D) and anti-p24 28B7 (MediMabs) antibodies for an additional 45min at RT and analyzed by flow cytometry on a BD LSRII. In all experiments, uninfected CEM cells were included to set the threshold of positivity. The detailed protocol of the HIV-Flow procedure can be found here: dx.doi.org/10.17504/protocols.io.w4efgte.

### HIV-RNA-FLOW-FISH

HIV latently infected ACH2 cells and uninfected CEMx174 cells were grown in separate flasks at a concentration of 0.5 million cells per mL in RPMI 1640 medium (Gibco, Life Technologies) supplemented with penicillin/streptomycin (Gibco, Life Technologies and 10% FBS (Seradigm). ACH2 cells were stimulated for 24 hours with PMA (50ng/mL, Sigma-Aldrich) and ionomycin (0.5 μg/mL, Sigma-Aldrich) prior to spiking into uninfected CEMx174 cells at different ratios obtained by serial dilution, starting with a highest frequency of 1,500 reactivated ACH2 cells per million CEMx174 cells. HIV+ events were identified using the HIVRNA/Gag assay as previously described [13,24]. Briefly, cells were stained with Fixable Viability Dye (eBioscience), anti-Gag KC57 (Beckman Coulter) by intracellular staining and labelled for HIV gag mRNA using the PrimeFlow RNA assay (ThermoFisher) before acquisition on a flow cytometer (FACS Fortessa, BD). Analysis was performed using FlowJo version 10 for Mac (Treestar). HIV+ cells were identified as cells co-expressing both HIV Gag protein and gag mRNA.

### Ultrasensitive p24 immunoassay

p24 in cell lysates was measured according to the method described previously [14] with some modification. In brief, prior to the p24 SIMOA, cell pellets were lysed at 4 ×10^6^ cells/ml with a cell lysis buffer containing 1% Triton x-100 in 0.5% casein (in PBS) and 50% Hi-FBS for 15min at room temperature, then frozen at −80°C until analysis. 35 μL aliquots of M-280 beads (Invitrogen/Life technologies, Cat#11206D) in microcentrifuge tubes were washed once with 1ml 1%BSA/PBS, then 3.5μL of 1 mg/ml normal mouse IgG (mIgG) (GenScript, Cat# A01007) was added. The frozen cell lysate was thawed in the 37°C water bath and centrifuged for 10min at 14,000 rpm at 4°C, and 0.3ml cell lysate supernatant was collected after spinning and added into the microcentrifuge tubes containing the washed beads and mIgG. Diluted samples using the cell lysis buffer were treated similarly. The tubes were incubated at 4°C for 3h with 360° rotation using a HulaMixer. The samples were subsequently centrifuged for 10min at 14,000 rpm at 4°C and the supernatant was collected and run on a HD-1 Quanterix Analyzer.

## Acknowledgments

The authors thank the US National Institutes of Health and the Canadian Institutes of Health Research for support (NIH R01 AI110173, R01 AI120698, R21 AI136731, R01 AI134406, R56 AI145407; CIHR 152977 and 364408) and Colleen O’Neill for assistance with preparation of the manuscript. D.E.K is a FRQS Merit Research Scholar. NC is supported by Research Scholar Career Awards of the Quebec Health Research Fund (FRQS, #253292)

